# Structural features underlying the activity of benzimidazole derivatives that target phosphopeptide recognition by the tandem BRCT domain of the BRCA1 protein

**DOI:** 10.1101/555623

**Authors:** Vadiraj Kurdekar, Saranya Giridharan, Jasti Subbarao, Mamatha B. Nijaguna, Jayaprakash Periasamy, Sanjana Boggaram, Kavitha Bharatham, Vijay Potluri, Amol V. Shivange, Gayathri Sadasivam, Muralidhara Padigaru, Ashok R. Venkitaraman

## Abstract

The tandem BRCT (tBRCT) domains of BRCA1 engage pSer-containing motifs in target proteins to propagate intracellular signals initiated by DNA damage, thereby controlling cell cycle arrest and DNA repair. Recently, we identified Bractoppin, a benzimidazole that represents a first selective small molecule inhibitor of phosphopeptide recognition by the BRCA1 tBRCT domains, which selectively interrupts BRCA1-mediated cellular responses evoked by DNA damage. Here, we combine structure-guided chemical elaboration, protein mutagenesis and cellular assays to define the structural features that underlie the biochemical and cellular activities of Bractoppin. Bractoppin fails to bind mutant forms of BRCA1 tBRCT bearing single residue substitutions that alter K1702, a key residue mediating phosphopeptide recognition (K1702A), or alter hydrophobic residues (F1662R or L1701K) that adjoin the pSer-recognition site. However, mutation of BRCA1 tBRCT residue M1775R, which engages the Phe residue in the consensus phosphopeptide motif pSer-X-X-Phe, does not affect Bractoppin binding. Collectively, these findings confirm a binding mode for Bractoppin that blocks the phosphopeptide-binding site via structural features distinct from the substrate phosphopeptide. We explored these structural features through structure-guided chemical elaboration of Bractoppin, synthesizing analogs bearing modifications on the left and right hand side (LHS/RHS) of Bractoppin’s benzimidazole ring. Characterization of these analogs in biochemical assay reveal structural features underlying potency. Analogs where the LHS phenyl is replaced by cyanomethyl (2091) and 4-methoxyphenoxypropyl (2113) conceptualized from structure-guided strategies like GIST and dimer interface analysis expose the role of phenyl and isopropyl as critical hydrophobic anchors. Two Bractoppin analogs, 2088 and 2103 were effective in abrogating BRCA1 foci formation and inhibiting G2 arrest induced by irradiation of cells. Collectively, our findings reveal structural features underlying the biochemical and cellular activity of a novel benzimidazole inhibitor of phosphopeptide recognition by the BRCA1 tBRCT domain, providing fresh insights to guide the development of inhibitors that target the protein-protein interactions of this previously undrugged family of protein domains.

## INTRODUCTION

Intracellular signals triggered by DNA double strand breaks (DSBs) initiate homologous recombination (HR) pathway as part of the DNA damage response (DDR) to repair these breaks [1a,1b].

The small molecule inhibitors of HR pathway proteins provide huge therapeutic potential in selective disruption and can be used as a chemo-sensitizer and/or in combination therapy along with inhibitors/compounds directed towards compensatory mechanisms or pathways or PARP inhibitors [2]. Current inhibitors of DSB repair proteins in HR pathway mostly are ATP-competitive protein kinase inhibitors of ATM [3], ATR or CHK1 that suffer from lack of specificity leading to off-target toxicities [4], owing to the structural similarity of human kinase domain families, and thus limit therapeutic exposure and efficacy. Hence, targeting protein domains that are downstream of kinase signaling pathway propagating signals via recognition of phosphorylated protein substrates could be an alternate strategy to overcome kinase mediated pleiotropic effects. The BRCT (BRCA1 C-terminal) domain originally identified in Breast cancer associated 1 (BRCA1) protein coordinates multiple signals of DDR by engaging phosphorylated proteins like BACH1[5], ABRAXAS[6], CTIP[7], ACC. Inhibition of BRCA1-ABRAXAS interaction functionally mimics BRCA1 tBRCT mutations leading to genome instability [8]. Further, it is observed that truncation or missense mutations in tBRCT domain of BRCA1 protein leads to increased risk for breast and ovarian cancers [9]. BRCA1 BRCT mutations M1775R and K1702M which disrupt phosphopeptide interactions are found to increase cell susceptibility for radiation damage and accumulation of cells at S/G2 phase of the cell cycle [10]. Our recent discovery of Bractroppin, the first small molecule inhibitor of BRCA1 tBRCT, inhibits BRCA1 recruitment to site of DNA breaks, and suppresses damage-induced G2 arrest and assembly of the recombinase, RAD51 without affecting MDC1 and TopBP1 recruitment [11]. These results provide proof of concept in using small molecule inhibitors against proteins involved in the transmission of the signals emanated from kinases in HR repair pathway.

In human BRCA1, two BRCT domains are arranged in a head-to-tail orientation, tandem BRCT (tBRCT) domains[12a,12b] and recognize phosphorylated substrates with the consensus sequence, pSer-X-X-Phe [13]. Structural analysis of the complex between BRCA1 tBRCT and BTB domain and CNC homolog 1 (BACH1), ABRAXAS, CTIP, ACC, reveals that the conserved pSer residue contacts the polar side chains of Ser1655 and Lys1702 from N-BRCT fold, the Phe residue engages a hydrophobic pocket formed by the side chains of Met1775, Phe1704, Arg1699, and Leu1839 from both N- and C-BRCT domains [15]. Natarajan and coworkers developed tetrapeptide (Ac-pSPTF-COOH) with 40 nM affinity and subsequently demonstrated a poly arginine containing cell permeable version - peptide 2 (Ac-R10G-pSPTF-CO2H) not only inhibits BRCA1-ABRAXAS/BACH1/CtIP interaction but also mimics BRCA1 mutations in cells. They later identified difluorophosphonate containing compound 15a with Kd 0.71 uM [13]. White *et al.* discovered non-phospho containing peptide inhibitor, Peptide 8.6 (Kd 3.6 uM) with glutamic acid as pSer mimic using mRNA display technology [14]. The peptidic nature and charged pSer limits their further development for therapeutic purpose therefore emphasizing the need for the development of Bractoppin as a small molecule inhibitor of BRCA1 tBRCT. Although, Bractoppin binding mode was validated by SAR, molecular level understanding of binding pocket residues responsible for activity will compensate for the lack of co-crystal structure and would give us impetus in building structure activity relationships for potency optimization. Here, we define the structural features that underlie the biochemical activities of Bractoppin by BRCA1 tBRCT pocket residue mutagenesis and apply multiple structure-guided chemical elaboration for potency optimization.

## RESULTS AND DISCUSSION

### Mutations that define the structural features responsible for Bractoppin’s activity

The consensus phosphopeptide motif pSer-X-X-Phe (numbered 0 to +3 repectively) binds at the interface of the tandem BRCT repeats of BRCA1 in such a way that the phosphopeptide’s pSer residue contacts the polar side chains of S1655 and K1702 of N-BRCT, while the +3 Phe residue engages a hydrophobic pocket formed by the side chains of M1775, F1704, R1699, and L1839 formed by both N- and C-BRCTs (Fig 1A). In our recent study, Bractoppin’s predicted binding mode on BRCA1 tBRCT was strengthened by Structure activity relationships (SAR) built by challenging the critical functional groups of Bractoppin predicted to be responsible for interacting with the pocket residue[11]. The predicted binding model of Bractoppin shows that Benzimidazole ring engages the pSer recognizing residues, the two hydrophobic groups, phenyl and benzyl explore two hydrophobic pockets that are not explored by the phosphopeptide (Fig 1B) [11]. Establishing the critical pocket residues by site directed mutational studies would pave the way for optimizing the potency of Bractoppin in a structure guided way. Hence, we chose four residues for mutational studies that affect either specifically peptide or compound binding or affect both (Fig 1A and 1B). i) F1662R mutation was selected based on peptide binding pocket analysis of structurally similar TOPBP1 tBRCT7/8 [16] where R1280 interacts with pSer/Thr of its cognate phosphopeptide. In BRCA1, the mutant F1662R protrudes into the hydrophobic pocket made up of F1662, L1657, V1654 and L1676 residues predicted to be occupied by phenyl ring of Bractoppin. The phenyl ring attached to 2nd position of Benzimidazole of Bractoppin on the left hand side (LHS) makes T-shaped, pi-pi stacking interaction with F1662. Thus, Arginine side chain would cause steric clash for Bractoppin binding while not affecting BACH1 phosphopeptide binding. ii) K1702 is a known critical residue for phosphopeptide recognition [17] and Bractoppin Benzimidazole core unsaturated nitrogen is predicted to make a H-bond interaction with K1702 side chain similar to that seen for pSer of phosphopeptide. Hence, K1702A mutation was predicted to abrogate both Bractoppin as well as BACH1 phosphopeptide binding. iii) M1775R is a well known cancer associated mutation, known to disrupt interaction with BACH1 phosphopeptide [15] at the hydrophobic core of BRCT - BRCT interface where +3 phenylalanine residue of phosphopeptide interacts with M1775 (PDB ID 1N5O) [17]. Based on the predicted binding mode, Bractoppin does not occupy this pocket, and hence it should not be affected by the mutation. iv) L1701K mutation is predicted to disrupt the hydrophobic groove between the two BRCT domains, thus precluding Bractoppin’s binding as the 2-Fluoro-benzyl group attached to the piperizine ring occupies it. The peptide binding should not be affected as there is no direct interaction, unless, the orientation of the two BRCT domains is affected.

**Figure 1.**
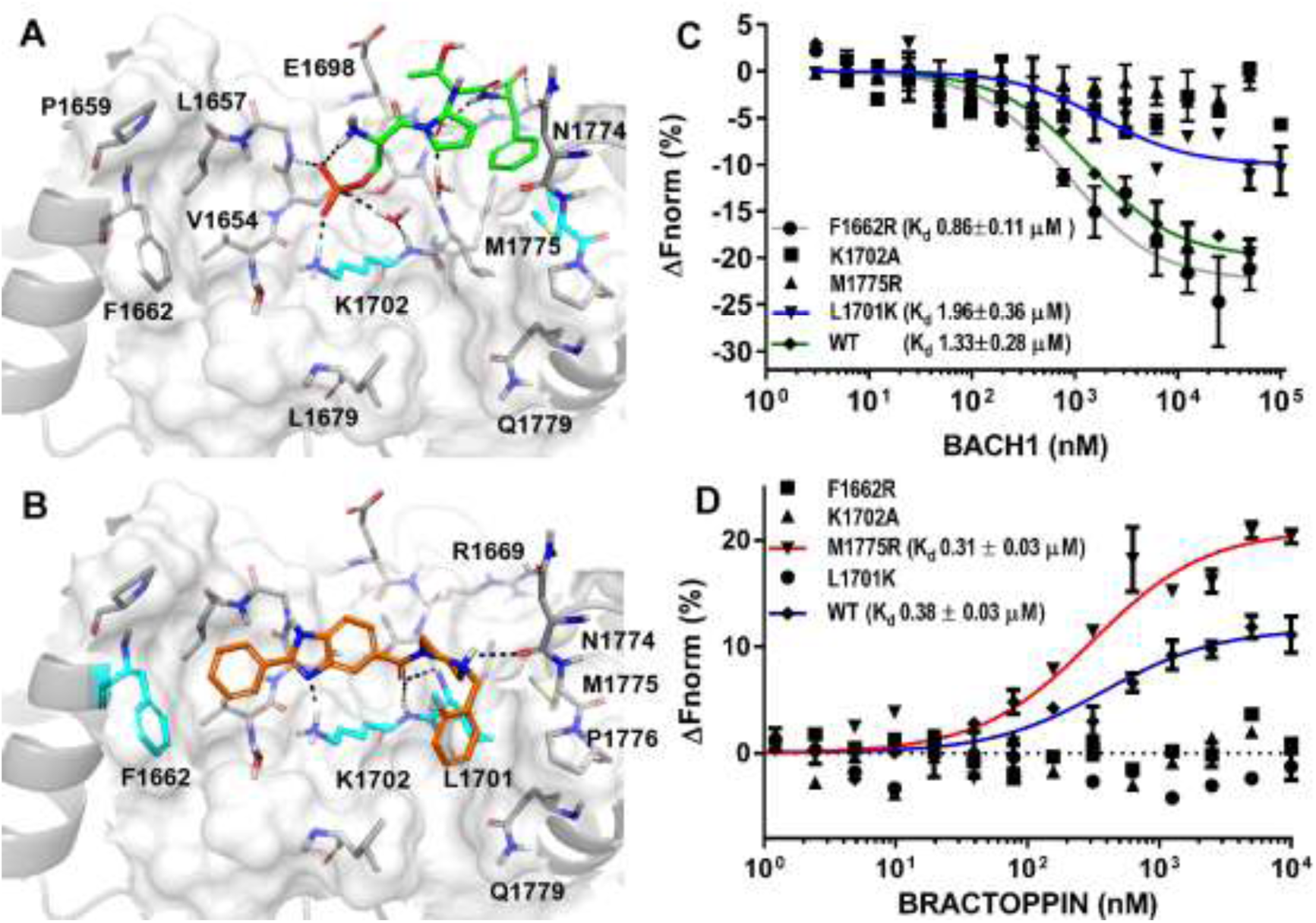
Bractoppin binding site validation by systematic mutations on BRCA1 tBRCT at common and distinct sites of interaction by Bractoppin and BACH1 phsophopeptide. Binding site on BRCA1-tBRCT (gray) of **A)** BACH1 phosphopeptide (green stick model) and B) Bractoppin (orange stick model). Residues selected for point mutations are marked in cyan color and Hydrogen bond interactions are represented as dotted line in black color. Direct binding plot of C) BACH1 phosphopeptide and D) Bractoppin with mutant and wild type BRCA1 tBRCT measured by MST. Titrant concentration in nM is plotted on the x axis, against changes in normalized fluorescence (ΔF_norm_), on the y axis. Plots represent the mean ± SD (error bars) from three independent experiments.

Experimental results obtained from direct binding Microscale Thermophoresis (MST) studies of BRCA1 tBRCT mutants with Bractoppin compared to its cognate phosphopeptide is in agreement with computational predictions. F1662R and L1701K mutants bind to BACH1 phosphopeptide with Kd of 0.86 uM and 1.96 uM respectively while the known M1775R and K1702A mutants have lost binding (ref) (**Fig. 1C**). In agreement with predicted (**Fig. 1B**) binding mode, loss of Bractoppin binding to K1702A mutant confirms that it competes with BACH1 phosphopeptide at phosphorecognition site. Bractoppin binds to wide type (WT) and M1775R mutant with similar affinity, Kd of 0.38 uM and 0.31 uM respectively, confirming that it does not occupy +3 phenylalanine binding pocket (Fig 1D). Further, Bractoppin occupies the two hydrophobic pockets as seen by the loss of its binding to F1662R and L1701K mutants. These mutational data together with previous SAR of critical functional groups responsible for Bractoppin’s binding define the structural features that are common and distinct in BRCA1 tBRCT responsible for Bractoppin’s binding from phosphopeptide. This implies that Bractoppin can bind to BRCA1 tBRCT wildtype and M1775R mutant, that are found in certain cancers and inhibit their phosphopeptide recognition. Further, it could be speculated that Bractoppin could preclude dimerization of BRCA1 tBRCT’s thus inhibiting diphosphorylated ABRAXAS. With a validated binding site of Bractoppin, we explored SAR at two positions, LHS of Benzimidazole group and RHS of the piperizine ring to improve potency.

**Scheme 1.**
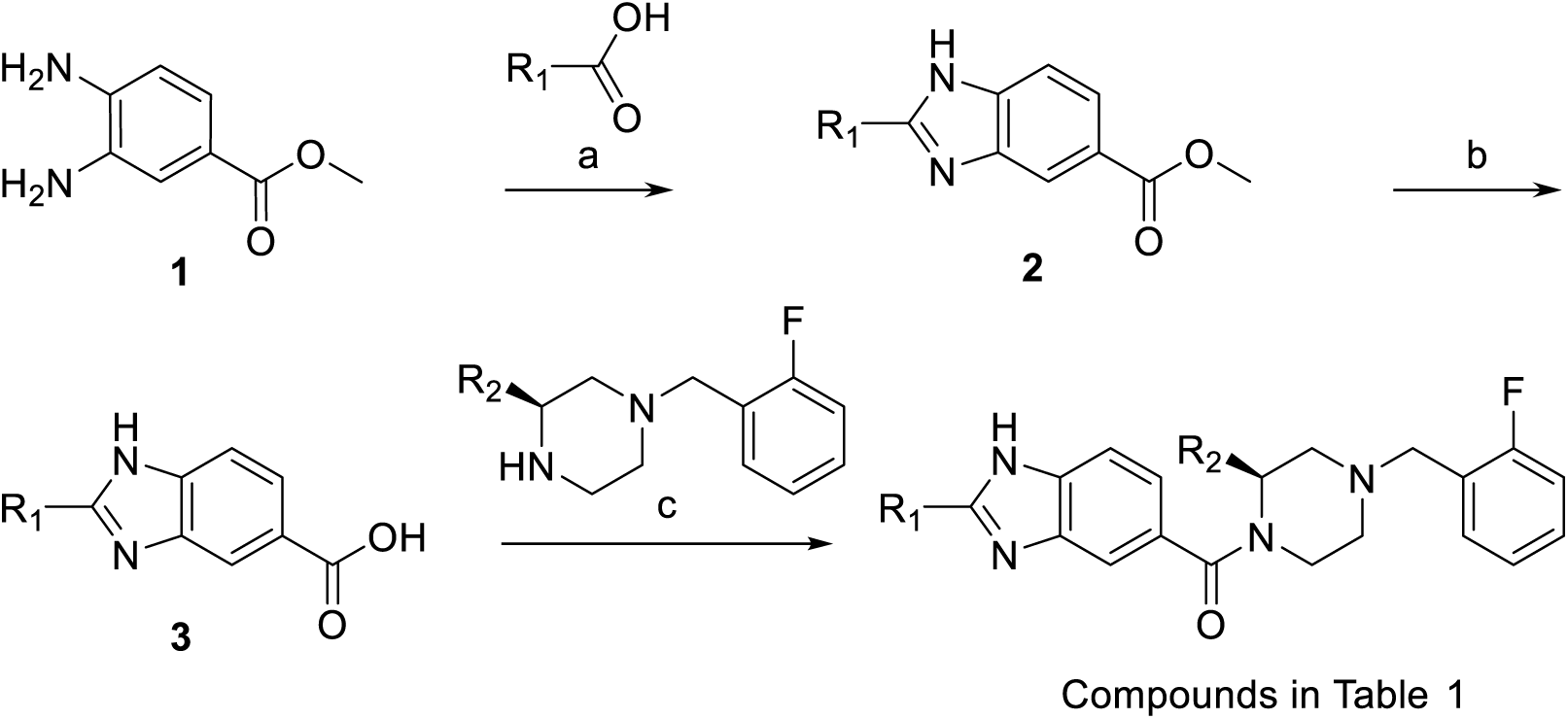
Reagents and conditions: (a) PPA, 170 °C; (b) AcOH, Conc. HCl 90 °C; (c) HATU, DIPEA.

### Structure based optimization of Bractoppin on the LHS

The phenyl ring on the LHS of Benzimidazole ring of Bractoppin occupies a hydrophobic pocket formed by F1662, L1657, V1654 and L1676 residues. To probe the pocket flexibility around the phenyl ring of Bractoppin, compounds with various modifications on the LHS (Table 1) were synthesized as shown in Scheme 1. The structure activity relationships of compounds were based on the competitive MST where the compounds competed with BACH1 phosphopeptide for BRCA1 tBRCT. The addition of 3,5-dimethoxy substitution on the phenyl ring (2010) decreased its activity by ∼ 9 fold. Compounds like 2090 with para methoxy phenyl, 2119 with para methoxyethoxy phenyl, 2088 with para cyanomethoxy phenyl had decreased activity by 3 fold, 15 fold and 3 fold respectively. Further, when the phenyl ring is replaced with 1,3,4-Oxadiazole (2086), it turned inactive. These data expose the role of phenyl as a hydrophobic anchor and restricts the orientation of the Benzimidazole ring optimal for binding. To further explore SAR on LHS, two independent strategies were used to propose modifications on Bractoppin: i) Thermodynamic analysis of water molecules at peptide binding site using GIST [18] ii) Analysis of the co-crystal structure of BRCA1 tBRCT dimer interface bound to diphosphorylated peptide of ABRAXAS.

**Table 1.**
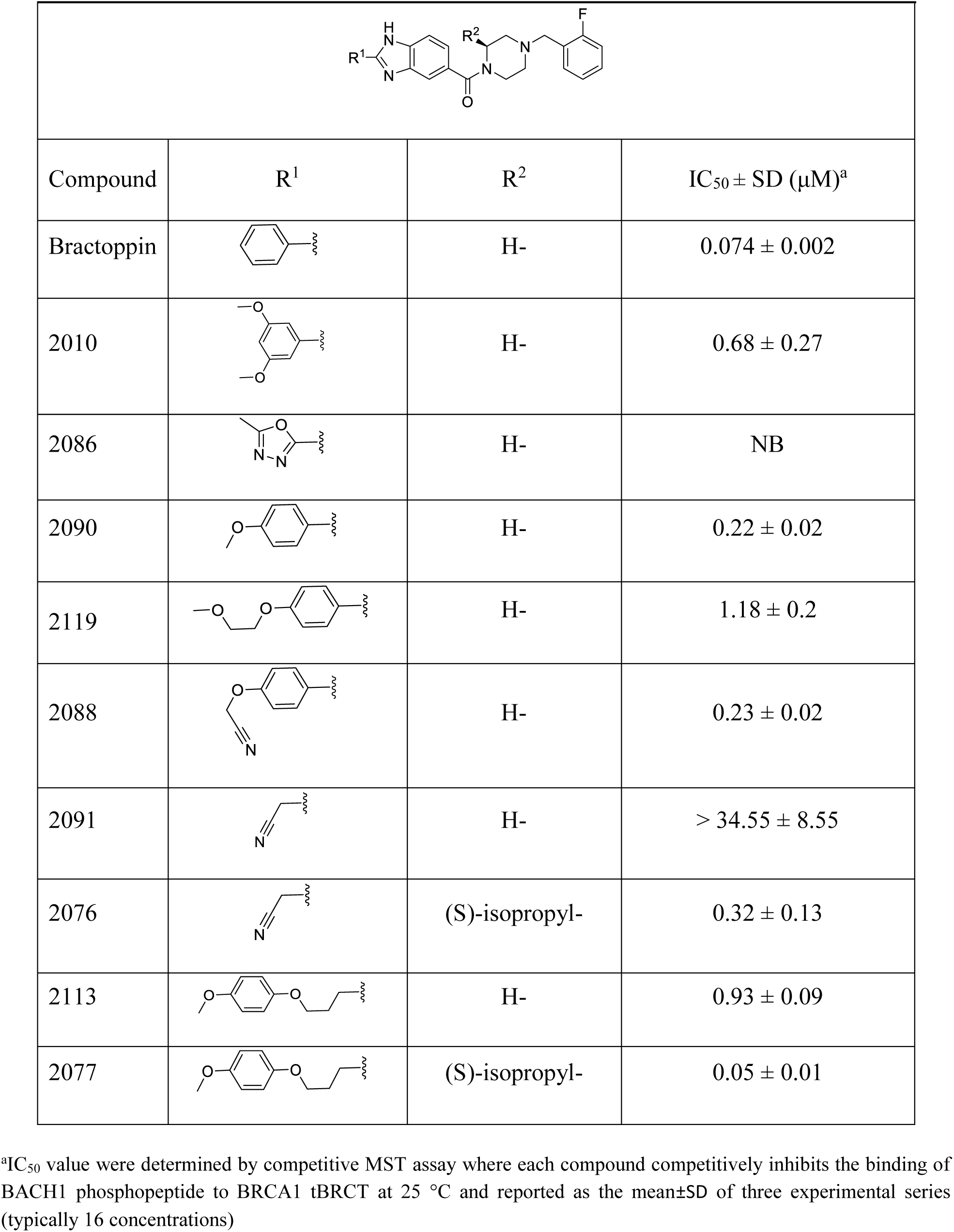
Structure-guided modifications on LHS at 2^nd^ position of Benzimidazole scaffold of Bractoppin.

|  |  |  |  |
| --- | --- | --- | --- |
| Compound | R<sup>1</sup> | R<sup>2</sup> | IC<sub>50</sub> ± SD (μM)<sup>a</sup> |
| Bractoppin |  | H- | 0.074 ± 0.002 |
| 2010 |  | H- | 0.68 ± 0.27 |
| 2086 |  | H- | NB |
| 2090 |  | H- | 0.22 ± 0.02 |
| 2119 |  | H- | 1.18 ± 0.2 |
| 2088 |  | H- | 0.23 ± 0.02 |
| 2091 |  | H- | > 34.55 ± 8.55 |
| 2076 |  | (S)-isopropyl- | 0.32 ± 0.13 |
| 2113 |  | H- | 0.93 ± 0.09 |
| 2077 |  | (S)-isopropyl- | 0.05 ± 0.01 |

<sup>a</sup>IC<sub>50</sub> value were determined by competitive MST assay where each compound competitively inhibits the binding of BACH1 phosphopeptide to BRCA1 tBRCT at 25 °C and reported as the mean±SD of three experimental series (typically 16 concentrations)

**Table 2.** Structure-guided modifications on RHS of the piperizine ring of Bractoppin

| Compound | X | R<sup>3</sup> | IC<sub>50</sub> ± SD (μM)<sup>a</sup> |
| --- | --- | --- | --- |
| 2048 | N | -CH<sub>3</sub> | NB |
| 2098 | N |  | 0.35±0.08 |
| 2093 | N |  | 0.97±0.2 |
| 2096 | N |  | 0.55±0.07 |
| 2103 | N |  | 0.02±0.003 |
| 2104 | N |  | 0.28±0.1 |
| 2099 | C |  | 0.05±0.01 |

<sup>a</sup>IC<sub>50</sub> value were determined by competitive MST assay where each compound competitively inhibits the binding of BACH1 phosphopeptide to BRCA1 tBRCT at 25 °C and reported as the mean±SD of three experimental series (typically 16 concentrations)

#### i)Water dynamics at peptide Binding site

GIST analysis highlights hydration sites at the peptide-binding site, suggesting most stable water densities along with their hydrogen directionality. Thermodynamic properties of hydration can be decomposed into enthalpic and entropic energies (supplementary Table S1). Water-entropy can be further decomposed into translational (-*TΔS*^*trans*^) and orientational (-*TΔS*^*orient*^) contributions and water-enthalpy as solute–water interactions (*ΔE*_*sw*_) and water–water interactions (*ΔE*_*ww*_). Conventionally we know that water molecules near to hydrophobic site are entropically less stable. Therefore, in conventional view we consider release of such unfavorable water to the bulk enhances ligand binding affinity [19]. Study of stable waters at pockets has gained significant attention in recent years to design potent inhibitors and to understand binding mechanism [20]. Analysis of 23 BRCA1 tBRCT crystal structures deposited in Protein Data Bank (www.rcsb.org) at the peptide binding site showed two possible rotomeric states of F1662. Therefore, to account for side chain flexibility, MD simulations were performed by constraining only protein backbone and Cα atoms of apo BRCA1 tBRCT unlike the conventional way, where all the protein atoms are constrained.

The stable water densities notated as w1 to w6 (Fig 2A) from GIST analysis at the site of interest, overlap with crystal waters except for w4 in phosphopeptide bound co-crystal structure, PDB ID 3K0K [25]. Water molecule, w1, held between backbone amine of V1654 and carbonyl oxygen of T1677 is highly stabilized by −11.78 kcal/mol energy. Since it is involved in the stabilization of protein, its displacement is predicted to be unfavorable by small molecule. Among the ordered network of waters, w3, w4, w5, w6, corresponding to the crystal waters in the Apo form (PDB ID: 1JNX) [17], peptide’s phospho group replaces w5 while preserving the rest of the water network bridging protein and peptide through H-bond interactions. Bractoppin backbone carbonyl oxygen displaces water, w3, explaining the crucial role of carbonyl group in binding as the removal of carbonyl renders the molecule inactive as reported in our previous publication (CCBT2908) [11]. Water w2 stabilized (−1.71 kcal/mol) by water-water interaction mainly with w1 is ideal for replacement. Hence, small molecules that displace w2 and make hydrogen bonding interaction with w1 will be entropically favored for binding. The w2 represented as electrostatic field points (Fig 2B) was utilized to grow Benzimidazole scaffold to identify new functional groups, which could mimic the interactions. Cresset Spark program was employed to search for modifications with similar features [19]. Compounds with Spark field point score more than 0.9 were considered and based on synthetic feasibility, compound 2091 was chosen as it was predicted to replace w2 and interact with w1 water. Contrary to our expectation, 2091 was found to be less active than Bractoppin with IC50 > 28 uM. The loss of potency could be due to the loss of phenyl, a critical anchor that could not be compensated by replacing w2 water and additional interaction with w1 water. However, addition of S-isopropyl on the piperizine ring in compound 2076 had not only recovered but has improved the potency to 0.32 uM (Fig 2C). This could be explained by the isopropyl group acting as a hydrophobic anchor wherein it protrudes towards pocket formed by L1701 and M1775 side chains [11].

**Figure 2.**
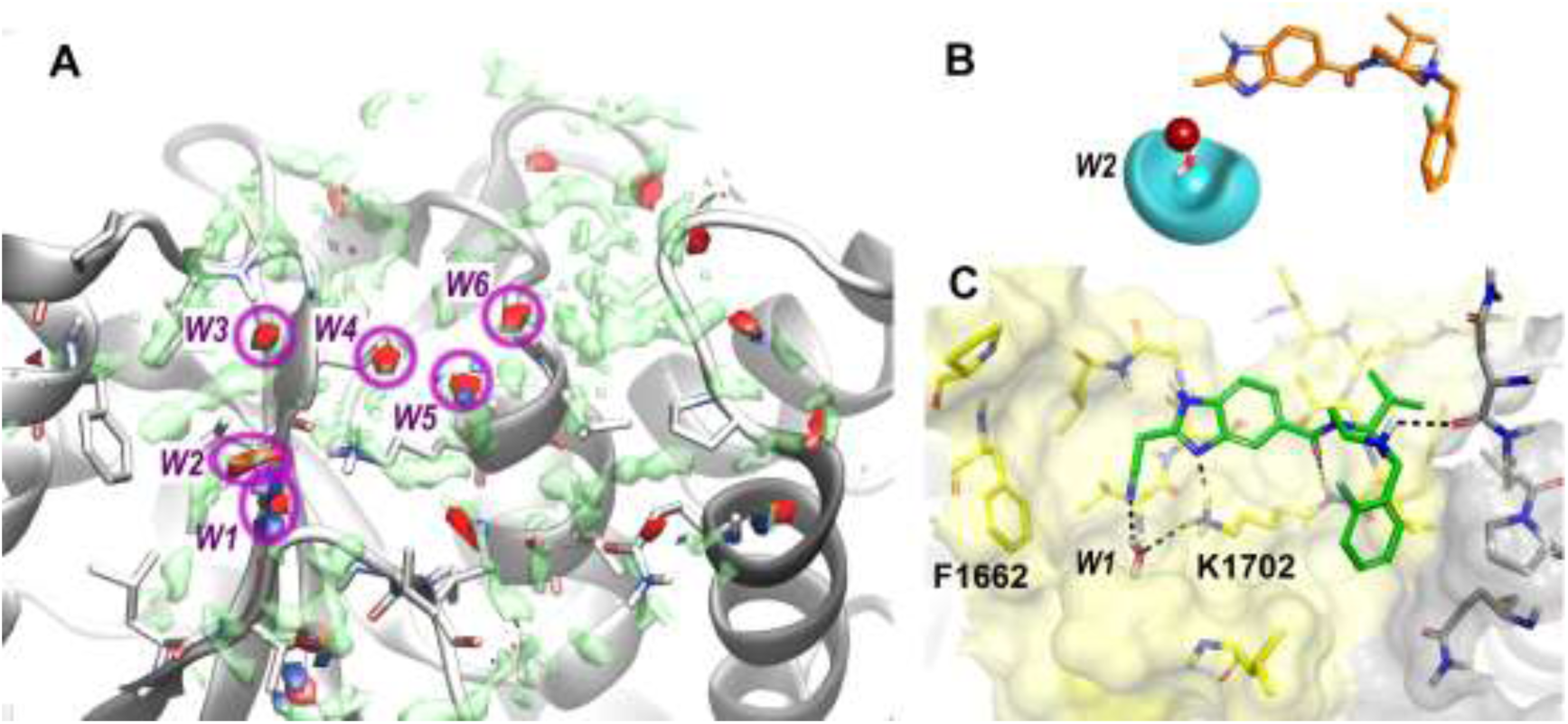
Structure-guided strategies to explore SAR around Bractoppin on the LHS of Benzimidazole ring using GIST analysis. **A)** Stable waters at ligand binding site of BRCA1 tBRCT with density of water oxygen (red) and hydrogen (blue) contoured at 10. Water density with high energy (Etotal > −0.25 kcal/mol) is shown as green isosurface. **B)** Field points used to search for new fragments to be attached at 2nd position (R1) of benzimidazole scaffold (orange stick model). Cyan contour and sphere represents negative field, yellow contour represents hydrophobic field and red spheres indicates positive field. Size of the spheres indicates strength of the interaction field. **C)** Binding modes of 2076 compounds shown as sticks (green). Protein is shown as surface representation with the two BRCT domains colored in yellow and grey. w2 water is shown in sticks and the H-bond interactions are shown in black dotted lines.

#### ii) Insights from the co-crystal structure of BRCA1 tBRCT dimer interface

The co-crystal structure of dimerized BRCA1 tBRCTs with double-phosphorylated ABRAXAS peptide (PDB 4Y18) [21] provided us a valuable starting point to look for contacts near the peptide binding site for optimization of Bractoppin. Structural analysis of the dimer interface revealed pi-pi stacking interaction between F1662 of chain A with Y1666 of chain B (Fig 3A). These structural features from Benzene ring of Tyr1666 were represented as electrostatic and steric field points (Fig 3B) using Spark to grow Benzimidazole scaffold by linking fragments/new functional groups. Among the fragments that mimic the interaction with Spark field point score of greater than 0.9 and considering synthetic feasibility, compound 2113 was synthesized which showed an activity of IC50 0.93 uM. However, addition of S-isopropyl on piperizine ring as in case of 2077 improved the activity with an IC50 of 0.05 uM (Fig 3C). This is similar to the pattern seen earlier with compounds 2091 and 2076 and contrary to the pattern observed for Bractoppin and 2010, with phenyl group as an anchor on the LHS. Overall, the modifications on the LHS flip flop between potent and weak activity due to the need for one hydrophobic anchor between the two anchor points, phenyl and S-isopropyl.

**Figure 3:**
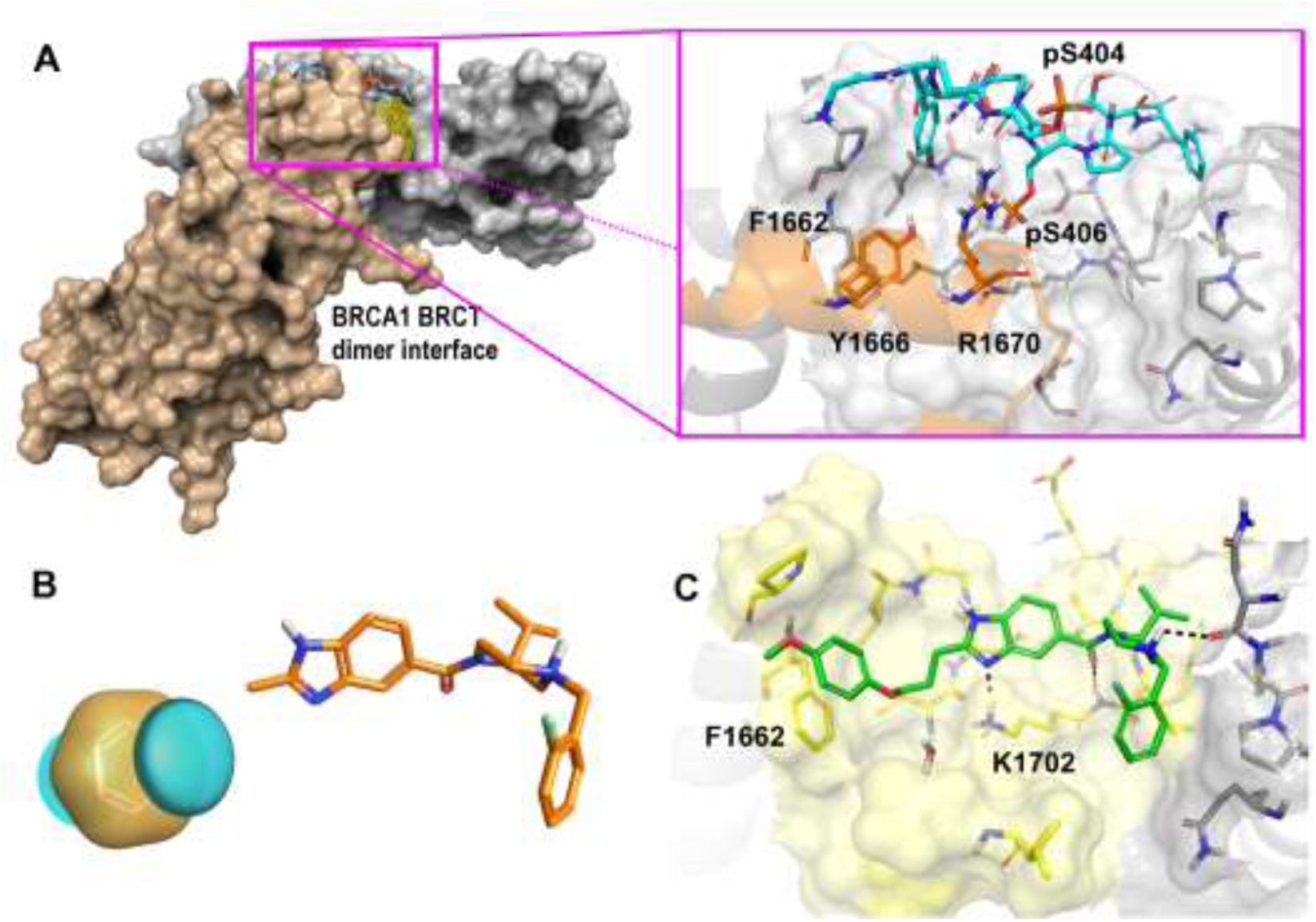
Structure-guided strategies to explore SAR around Bractoppin on the LHS of Benzimidazole ring using tBRCT dimer interface analysis. **A)** BRCA1 tBRCT dimer formed by diphosphorylated Abraxas[22] reported in PDB ID 4Y18. Gray surface view represent one tBRCT while light orange colored surface view represents another tBRCT involved in forming dimer, cyan stick model represent diphosphorylated peptide derived from ABRAXAS with zoomed view at dimer interface at peptide binding site. **B)** Field points used to search for new fragments to be attached at 2nd position (R1) of benzimidazole scaffold (orange stick model). Cyan contour and sphere represents negative field, yellow contour represents hydrophobic field. **C)** Binding mode of 2077 compounds shown as sticks (green). Protein is shown as surface representation with the two BRCT domains colored in yellow and grey. w2 water is shown in sticks and the H-bond interactions are shown in black dotted lines.

### Structure based optimization of Bractoppin on the RHS

On the RHS, based on the MD simulations analysis of BRCA1 tBRCT and Bractoppin complex structure, we explored interactions that could be captured at the interface of tBRCTs. The hydrophobic groove at the interface of two BRCT domains has P1776, and L1701, N1774 Q1779 and L1679 (Fig 1B). The compounds with various modifications on the RHS were synthesized as shown in Scheme 2. It was clear from our earlier results that the para position of benzyl could tolerate methyl (2107, IC50 = 0.3 uM) but not hydroxyl (2106, IC50 = 3.6 uM) indicating that the para-position is masked in the hydrophobic pocket [11]. To gauge the significance of o-Fluro benzyl, we replaced it with methyl as in 2048 and extended the benzyl ring with homo benzyl as in 2098. Complete loss of activity of 2048 shows the importance of RHS hydrophobic group. Reduction in activity by 5 fold for 2098 compared to Bractoppin corroborates with loss of activity for 2107 [11]. The ortho position substituents, 2093, the defluoro version turned out to be as potent as Bractoppin and the o-methoxy substitution (2096) had 7 fold decreased potency. Substitution at the meta position with morpholino group (2103), has affinity similar to Bractoppin. The binding model shows that the morpholino is solvent exposed suggesting that this position could be ideal for further optimization of solubility without affecting the affinity of the compound. The possibility of Q1779 to flip as seen in the available crystal structures has motivated us to check if compound 2104 could capture H-bond interactions. But, decrease in potency by 5 fold suggests that the flipped rotamer may not be available for interaction. Further, to enhance the scope of chemistry on the piperizine ring, it was changed to piperidine ring in 2099. From the MD simulations of Bractoppin suggest that the protonated nitrogen’s H-bond interaction with N1774 shown in dock pose is not essential as it is lost during the simulations (Supplementary Fig S1). As predicted, 2099 retained potency and has opened new avenues for chemistry. Overall, there were modest improvement in potency with modifications on RHS and LHS.

**Scheme 2:**
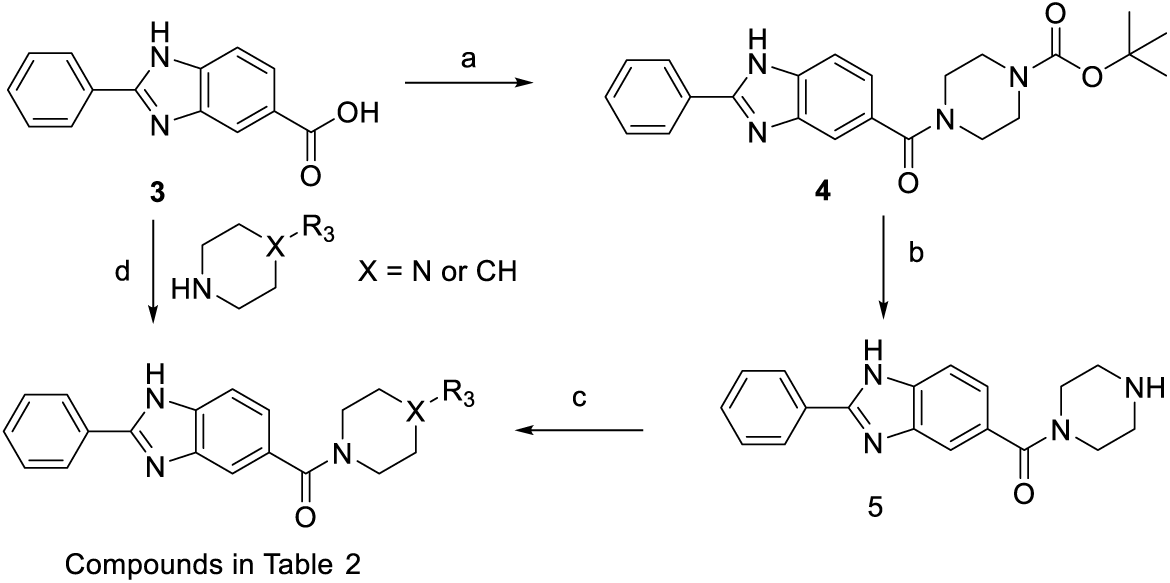
Reagents and conditions: (a) tert-butyl piperazine-1-carboxylate, HATU, DIPEA, DMF; (b) 4N HCl.Dioxane; (c) Na(CH_3_COO)_3_BH, AcOH, DCM; (d) EDC, HOBt, DMF.

### Cellular effects of potent Bractoppin analogues

Upon irradiation (IR), BRCA1 protein assembles as microscopic foci at sites of double strand DNA breaks through phosphorylated ABRAXAS via the BRCA1 tBRCT domain [6,22]. Overexpression of BRCA1 tBRCT domain, competes for the phosphorylated substrate binding to the full length endogenous BRCA1 protein, rendering it inactive and is used as a positive control for foci inhibition in the assay [11]. Soluble compounds with RHS modifications (2103); LHS modifications (2088); Bractoppin and its negative control, 2048, were tested for their inhibitory potential in comparison with the BRCA1 tBRCT overexpression phenotype. Significant inhibition of BRCA1 foci recruitment was observed with both compounds 2103, 2088 similar to Bractoppin but not with its inactive analogue, 2048 (**Fig 4**). Indeed, compounds that inhibited BRCA1 foci recruitment also abrogated the G2 checkpoint [11] since the accumulation of BRCA1 at sites of DNA damage initiates events that leads to cell cycle arrest at G2 [23,24]. Together these finding demonstrate that 2088 and 2103 interrupt intracellular signaling that is essential for the control of DNA damage in cells.

**Figure 4:**
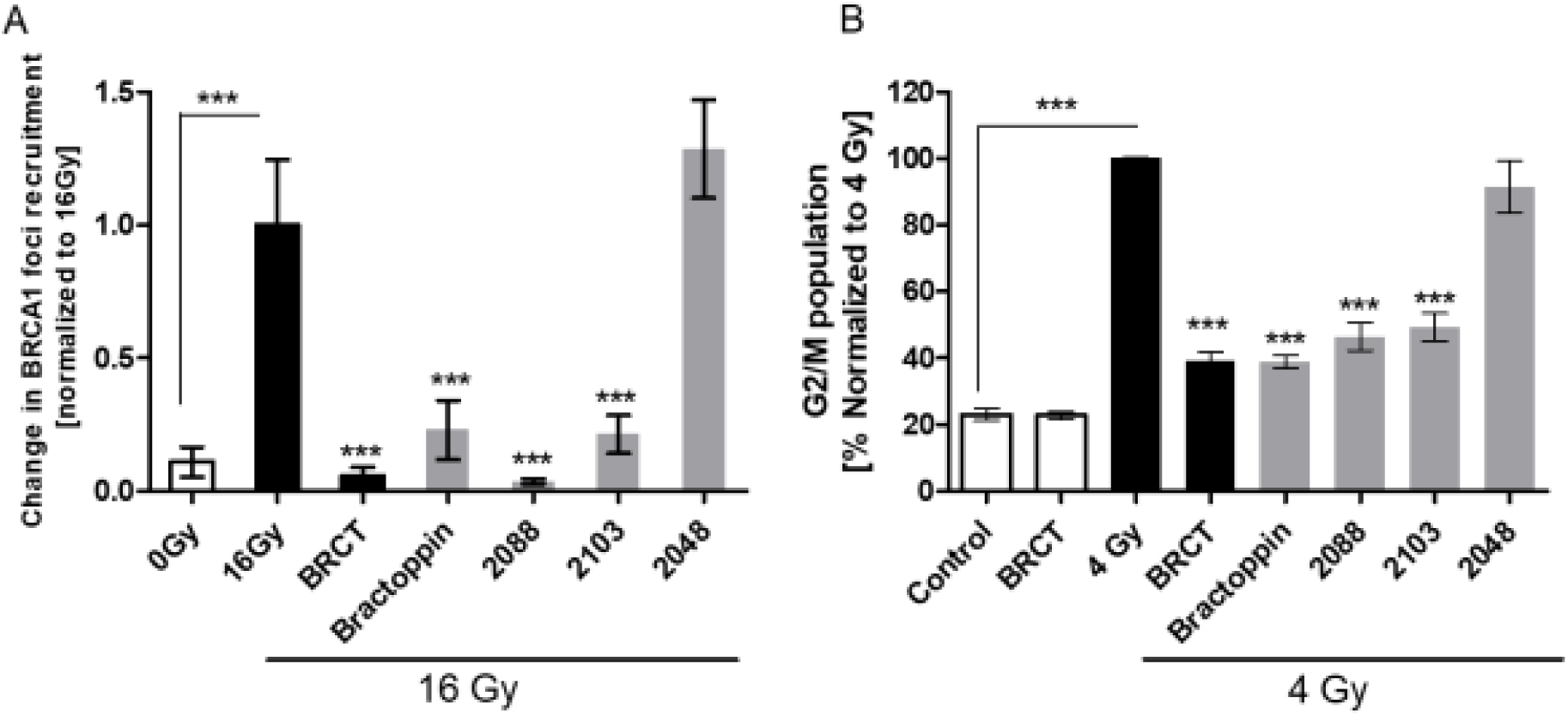
Bractoppin and its analogues disrupt intracellular signaling that controls DNA damage in cells via the tBRCT domains. **A)** Quantitative measurement of BRCA1 protein recruitment into nuclear foci at sites of DNA damage upon irradiation. Conditions include (untreated cells [0 Gy]; irradiation alone [16 Gy]; Tet-induced BRCA1 tBRCT expression 30 hr before irradiation; 100 μM Bractoppin or its analogues added 6 hr after irradiation. Data is normalized to 16Gy irradiation and calculated for change in cells positive for radiation-induced nuclear BRCA1 foci (mean ± SD; n = 15,140, 0 Gy; 11,098, 16 Gy; 11,175, BRCA1 tBRCT; 18,227, Bractoppin; 11,911, 2103; 6,266, 2088 and 16,063, 2048) enumerated by high-content imaging (for details see methods). Statistical significance was determined using an unpaired two-tailed t test. ***p ≤ 0.001. Similar results were observed in three independent repeats. **B)** Percentage of cells with 4N DNA content at the G2/M phase of the cell cycle by flow cytometry following DAPI staining. Cells were irradiated with 4 Gy at 8 hr after synchronous release into the cell cycle from thymidine block, and measurements made 16 hr later. Treatment conditions include, unirradiated cells (Control), or cells exposed to 4 Gy, with or without additional treatments using Tet-inducible BRCA1 tBRCT expression, 100 μM Bractoppin or its analogues. Tet-induced BRCA1 tBRCT expression was for 32 hr before radiation, while compounds were added 0.5 hr before. A total of 15,000 cells were analyzed per condition and data normalized to 4Gy irradiation, in replicates of 3. Results are representative of three independent experiment. Statistical significance was tested using an unpaired, two tailed t-test. ** P≤0.001

## CONCLUSIONS

Defining the critical residues responsible for Bractoppin’s activity at the phosphopeptide recognizing pocket of BRCA1 tBRCT will compensate for the lack of a co-crystal structure in understanding the molecular level interactions for building structure activity relationships and potency optimization. Residues that were predicted to make significant contribution in binding were mutated to tease apart the common and distinct binding modes of phosphopeptide and Bractoppin. Mutation of hydrophobic single pocket residues (F1662 and L1701) of BRCA1 tBRCT that do not interact with phosphopeptide present on either side of pSer recognizing residues abrogate Bractoppin binding but not BACH1 phosphopeptide. However, M1775R mutation which abolishes BACH1 binding has no impact on Bractoppin binding. These single residue mutations established the proposed molecular inteactions made by binding mode where it occupies the phospho recognizing site and two hydrophobic pockets of BRCA1 tBRCT that are common and distinct from peptide binding respectively. Binding of Bractoppin to M1775R germ line mutant can provide new insights to characterize functional impact and potentially expand its scope to probe its effect in cancers with BRCA1 M1775R mutation. After modifications on the meta and para positions of the LHS phenyl ring led to decrease in potency, structure guided strategies like GIST and dimer interface analysis were applied to explore SAR. Compounds were synthesized to validate the hypothesis of 2091 capturing a H-bond interaction with stable water, w2 and 2113 to capture T-shaped pi-pi stacking interaction with F1662. The loss of phenyl anchor effected their potency but was recovered by the addition of isopropyl anchor on the piperizine ring. This implies that Bractoppin series compounds depend on either of a hydrophobic anchor along with phospho site interaction made by Benzimidazole ring similar to the phosphopeptide. Moreover, Bractoppin and 2113 can abrogate BRCA1 tBRCT dimerization induced by diphosphorylated ABRAXAS. On the RHS, we could conclude that one plane of the o-Fluoro benzyl group was solvent exposed and meta position was optimal for adding groups without affecting activity. Bractoppin analogues, 2088 and 2103, each from LHS and RHS modifications was affective at 100 uM in abrogating BRCA1 foci and inhibiting G2 arrest upon irradiation in cells. Our work emphasizes the significance of combining multiple structure-guided strategies like pocket mutations, GIST and dimer interface analysis in proposing testable hypothesis that would lead to the development of inhibitors specifically for undrugged protein-protein interactions.

## MATERIAL AND METHODS

### Compounds Synthesis

Compounds were synthesized by O2h Discovery (Ahmedabad, India), and characterized by 1H-NMR and liquid chromatography coupled to mass spectrometry (LC/MS) and. Synthetic methods are provided in supporting material. All compounds were >95% pure as determined by high-performance liquid chromatography (HPLC). Stock solutions were prepared from dry powder in 100% DMSO at 50mM concentration. For MST assays, stock solutions were diluted to the indicated assay concentrations in 2% DMSO. For cell-based experiments, 20mM stocks of Bractoppin and its analogues in 100% DMSO were diluted in growth media (DMEM supplemented with 10%FBS, 2mM Glutamine) to the indicated concentrations (0.5% DMSO final concentration), thoroughly mixed, and spun at 13,000 rpm for 10 seconds before use.

### Generation of mutant clones of BRCA1 tBRCT

Site-Directed Mutagenesis was used to create mutant proteins with F1662R, K1702A, M1775R and L1701K using Phusion polymerase (Thermo Scientific, MA, USA) and nucleotide primer pairs as shown in Table 3. The PCR was carried out with the mixture containing forward and reverse primer separately for 2 cycles of 10 seconds at 98°C and 30 seconds at 55°C, followed by 20 seconds/kb plasmid size at 72°C. These two mixtures were mixed and run for 14 cycles of 10 seconds at 98°C and 30 seconds at 55°C, followed by 20 seconds/kb plasmid size at 72°C. The amplified product was then treated with restriction enzyme DpnI for 1 h at 37 °C and transformed into *E. coli* DH5α and mutant plasmids were verified by DNA sequencing.

**Table 3:**
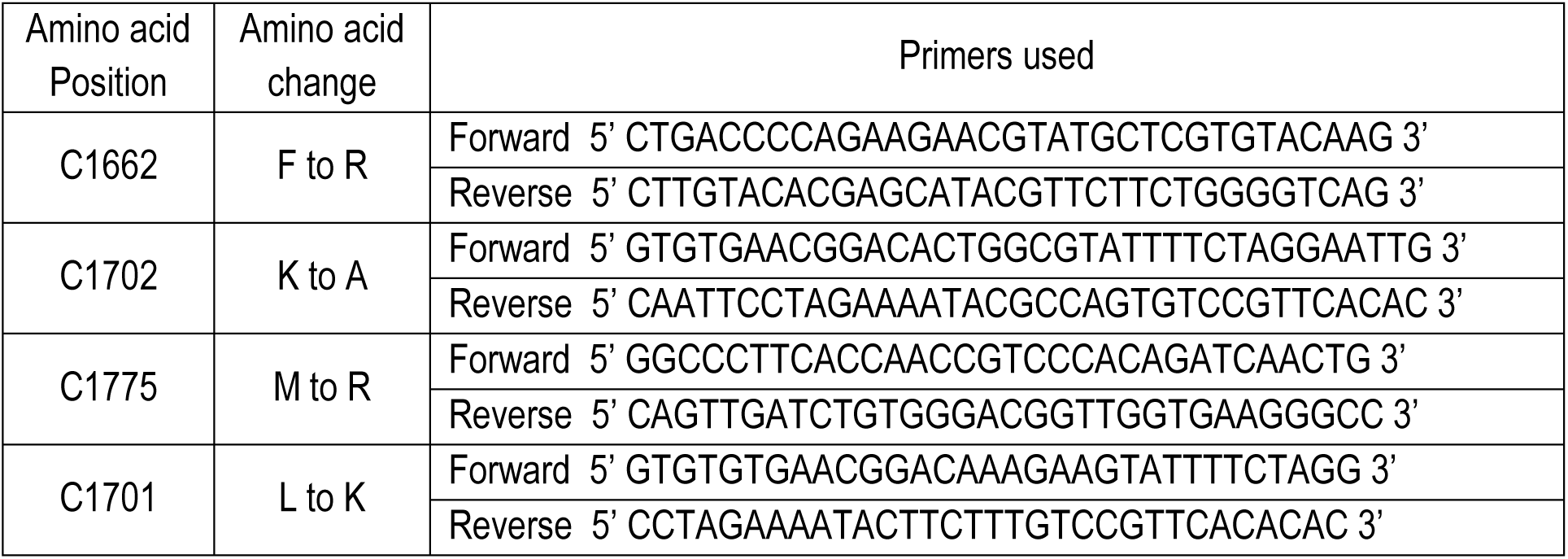
Primers used for site directed mutagenesis of BRCA1 tBRCT

### Protein expression and purification

The mutant clones generated as above was derived from **w**ild type synthetic gene construct encoding BRCT domain of BRCA1 tBRCT residues 1646-1859 with N-terminal 6x Histidine residues, and codon optimized for expression in *E. coli*, were procured (GeneArt, Regensburg, Germany) in the pET28a expression vector and expressed in *E. coli* cells in LB medium containing 50*µ*g/mL Kanamycin. 6x His-BRCA1 tBRCT expression was induced in BL21(DE3) strain at 0.6-0.8 OD_600_ with 0.25mM isopropyl β-D-1-thiogalactopyranoside (IPTG) at 18°C for 16 hours. Cells were harvested and the pellet was suspended in ice cold lysis buffer (50mM Tris HCl [pH 7.5], 400mM NaCl, 0.1mM PMSF, 1mM DTT, and 1 protease inhibitor tablet (Roche)). Cells were lysed by sonication on ice and centrifuged at 20,000 rev min^-1^ for 30 minutes at 4° C to remove cell debris. The supernatant was applied onto a HisTrap HP column (GE Healthcare) pre-equilibrated with a buffer (50mM Tris HCl [pH 7.5], 400mM NaCl, 1mM DTT, and 25mM Imidazole). The column was washed with same buffer until all unbound proteins were removed. The protein of interest was eluted using a linear gradient of 100% elution buffer (50mM Tris HCl [pH 7.5], 400mM NaCl, 1mM DTT, and 500mM Imidazole). Protein purity was visualized by running SDS-PAGE. Fractions of sufficient purity were pooled and concentrated to 2 ml using a 10 kDa cutoff Centricon centrifugal filter devices (Millipore). The concentrated protein was further purified using HiLoad 16/600 Superdex-75 prep-grade gel-filtration column (GE Healthcare) pre-equilibrated with 20mM Tris HCl [pH 7.5], 100mM NaCl and 1mM DTT.

### Microscale Thermophoresis (MST)

BRCA1 tBRCT wild type or mutant domains used in the assay were labeled with NT-647-NHS fluorescent dye using the Monolith NT™ Protein Labeling Kit (NanoTemper Technologies). Assays were carried out in 20mM Tris buffer, pH 7.4, with 200mM NaCl, 0.05% Tween-20 and 2mM DTT. Direct binding assay was carried out using 10μl of labeled protein at a final concentration of 20nM, mixed with 10μl of cognate peptide or test compound incubated on ice for 10 minutes followed by centrifugation at 15000 rpm at 4°C for 10 minutes. MST analysis was performed by loading 4µl of the supernatant into premium glass capillaries (NanoTemper Technologies) at MST power of 40% and LED power of 80%, at 22°C temperature using a Monolith NT.115 (NanoTemper Technologies). An initial “Capillary Scan” was performed to scan for fluorescence across the capillary tray to determine the exact position of each capillary before the MST measurement was started. Both peptide and test compounds were assayed at 16 different concentrations by serial dilution, and data were analysed using NanoTemper analysis software. Kd values were determined using “T-jump + Thermophoresis” settings. The change in thermophoresis between different experimental conditions was expressed as the change in the normalized fluorescence (ΔF_norm_), which is defined as F_hot_/F_cold_ (F-values correspond to average fluorescence values between defined areas in the curve under steady-state conditions under control (F_cold_) or experimental (F_hot_) conditions. Titration of the non-fluorescent ligand causes a gradual change in thermophoresis, which is plotted as ΔF_norm_ to yield a binding curve, which was then fitted to derive binding constants.

### Computational Methods

#### Protein Preparation

The crystal structure of BRCA1 BRCT in complex with minimal recognition motif (PDB ID: 3K0K) was retrieved from Protein Data Bank [25]. Using Schrödinger’s Protein Preparation Wizard (Schrödinger Release, 2015-3: LigPrep, 2015) protein structure was processed by adding hydrogen, fixing bond orders, fixing missing atoms and residues, removing ions, determining and fixing protonation states of side chains at pH 7.0. Protein was termini’s were capped with an N-terminal acetyl and a C-terminal amide group. The structure was then subjected to a restrained energy minimization step using the OPLS2005 force field with a maximum permitted RMSD of 0.30 Å

### Molecular dynamic (MD) simulations and analysis of water dynamics by Grid Inhomogeneous Solvation Theory (GIST)

Thermodynamics of water in ligand binding site was analyzed by Grid based Inhomogeneous Solvation Theory (GIST) developed by Nguyen and co-workers [18]. Change in the free energy at the hydration site ΔGSolv can be estimated by Equation (i) [26].

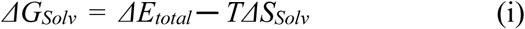

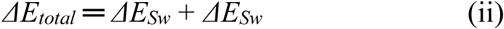

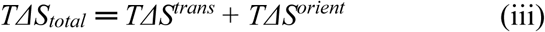

Where Δ*E*_total_accounts for change in solvation enthalpy and is calculated using Equation (ii). *ΔE*_*Sw*_accounts for change in solute-water interaction energy; *ΔE*_*ww*_accounts for change in water-water interaction; Δ*S*_total_accounts for change in solvation entropy at given temperature *T* can be calculated using Equation (iii). Where *TΔS*^*trans*^ accounts for translational entropy of water and *TΔS*^*orient*^ accounts for orientational entropy of water.

To calculate ΔGSolv at each grid points in defined area a molecular simulation was carried out in explicit water model using Amber15 MD simulation package [26,27] with ff14SB amber force field. BRCA1 BRCT domain PDB prepared earlier was used after stripping of water and bound phosphopeptide. Protein was solvated in rectangular box with TIP3P water molecules with minimum 12 Å from the protein surface. System was neutralized by counter ion Na+. Using periodic boundary conditions (PBC) and Particle Mesh Ewald method for long range electrostatic interaction calculation, system was minimized by 1500 steps with steepest descents algorithm and subsequently by conjugate gradient method for a maximum of 2000 steps keeping protein atoms constrained by 100 kcal/mol/Å. The system was gradually heated from 0 K to 50K for 20ps and then again gradually heated to 300K for 100 ps under NVT condition. Finally system was equilibrated under NPT condition for 2 nsat a constant pressure of 1 atm, followed by NVT condition for 1 ns. Finally 10 ns simulation was performed under NVT condition by keeping protein constrained at 100 kcal/mol/Å using pmemd.cuda [28] running on Nvidia K40 Tesla GPU card. During MD simulation all protein atoms were harmonically restrainedwith a force constant of 100 kcal/mol/Å. Particle Mesh Ewald method was used for long range electrostatic interaction calculation and a 9 Å cutoff was used to calculate all nonbonded interactions. 10000 frames obtained during 10ns simulation was used for GIST analysis. To account for possible rotamers of Phe1662, production run was repeated by harmonically restraining only protein backbone including Cα atom with a force constant of 100 kcal/mol/Å.

### Docking

To dock compounds, the bound peptide and water molecules were deleted except for 3^rd^ water molecule of Chain A, since it forms bridging interaction between backbone atoms of Val1654 and Thr1677. A grid box of size 22 x 27 x 22 Å3 with an inner box (10 x 15 x 10 Å3) centered at X, Y, Z coordinates −23.75, 48.00 and 4.0 was generated with default parameters and no constraints to cover proposed ligand binding site. Ligands were drawn and prepared using LigPrep. Ionization and tautomeric states were carefully selected after inspection. Five conformers were generated for each ligand using Confgen with an OPLS2005 forcefield and a minimum RMSD cutoff of 1.0 Å. Each conformer was then individually docked using the Glide SP protocol, to identify the 10 best poses per ligand. In case of Ligands designed by fragment swapping experiment performed by Cresset Spark [21], initially “refine only dock” was performed using Glide SP protocol followed by flexible docking protocol as mentioned earlier. Larger Deviation in interaction patterns observed in dock poses by both methods over Spark output pose was used as one of the filtering criteria.

### Expression Constructs

Tetracycline (Tet)-inducible plasmids encoding mCherry fused to wild-type BRCA1 tBRCT domains were prepared in the pcDNA5/FRT/TO-mCherry vector by gene synthesis (GeneArt, Regensburg, Germany). Synthetic polynucleotides encoding SV40-NLS (3X)-BRCA1 tBRCT (aa 1620-1862) were cloned between the BamH1 and XhoI restriction sites of the vector.

### Cell Lines and Cell Culture

The Flp-In™ T-REx™ 293 Cell Line was procured from Thermo Fisher Scientific (R78007) and maintained in DMEM supplemented with 10%FBS, 2mM L-glutamine, Blasticidin (15μg/ml) and Zeocin (100μg/ml) at 37°C with 5% CO2. Stable cell lines were generated by co-transfecting pOG44 with pCDNA5/FRT/TO vectors encoding BRCA1 tBRCT (aa 1620-1862) in a 9:1 ratio using FuGENE HD transfection reagent (E2311, Promega). After 10 days of selection using Blasticidin (15μg/ml) and Hygromycin B (50μg/ml), viable colonies were expanded, and assayed for the loss of β-galactosidase activity and Zeocin resistance to identify clones with stable integration of plasmid. Protein expression was induced with Doxycycline (1μg/ml for 48 h). Flp-In™ T-REx™ 293 cells were authenticated at the DNA Forensics Laboratory Pvt. Ltd., New Delhi, India using short tandem repeat (STR) profiling at the 8 core loci plus amelogenin specified by Capes-Davies et al. [29]. Flp-In™ T-REx™ 293 cells were confirmed to be female, and were an exact match (15/15 alleles) for the HEK-293 (CRL-1573) human cell line in the reference database.

### Cell Irradiation

Cells were irradiated to the indicated doses using either the Blood Irradiator 2000 with Cobalt-60 (Board of Radiation and Isotope Technology, Department of Atomic Energy, Government of India) at an effective dose rate of 3.9Gy/min, or with an X-ray generator (Xstrahl, RS225) at a dose rate of 1.5Gy/min.

### Immunofluorescence Staining for Damage-Induced Foci

HEK293 cells stably harboring plasmids for Tet-inducible expression of BRCA1 tBRCT domains were seeded at 30,000 cells/well on Matrigel-coated 96-well plates, and treated as indicated. Cells were fixed in 2.5% PFA, for 20’ at RT and incubated in 1x PBS with 10% FBS plus 0.5% TritonX-100 for 1 h for blocking and permeabilization. Primary antibody staining for endogenous BRCA1 was performed with mouse monoclonal Ab, sc-6954, Santa Cruz Biotech, (1:600 dilution) in PBST-BSA buffer (0.7mg/ml BSA, 0.05% Tween-20 in 1XPBS) for 1h at RT. Cells were then extensively washed in PBST-BSA buffer and stained with goat-anti-mouse Alexa 488 secondary antibody (Invitrogen), at 1:1000 dilution along with 2.5μg/ml DAPI for nuclear staining. Images were acquired on a high-content imaging platform (Cellomics ArrayScan VTI HCS Reader (Thermo Fisher Scientific) using a 40x objective. On average ∼800 fields from 6-well replicates, containing a total of ∼10-20K cells were imaged per treatment group and quantified using image analysis software (in-house algorithms using MatLab and commercially available HCS studio 2.0 from Thermo Fisher Scientific). Briefly, nuclear objects were defined, and foci were enumerated for the BRCA1 protein. Plots were generated to compare control (0 Gy) and 16 Gy irradiated samples for foci number vs. percentage of cells. Cut-off values for foci number per cell specifying the maximal difference between control and irradiated samples were determined to calculate the percentage of cells positive for radiation-induced foci. These cut-off values were used to enumerate changes in the percentage of cells positive for radiation-induced foci with or without inhibitor treatment.

### Cell Cycle Profiles

HEK293 cells stably harboring plasmids for Tet-inducible expression of tBRCT domains were seeded on 12-well plates at 1.8 X105 cells/well. Cells were thymidine blocked and synchronously released into the cell cycle following irradiation at 4Gy, without or with exposure to compounds (at final concentrations with 0.5% DMSO) in serum-containing media for 16 h. Cells were resuspended in 1x PBS, fixed with 80% ethanol, and permeabilized in buffer containing 0.1% Tween-20 in PBS (PBST), for 20’ at RT. DAPI staining was used to quantify nuclear DNA content. Analysis was performed using a Beckman Coulter Gallios analyzer, and quantified using the Dean-Jet algorithm in FlowJo software.

## ABBREVIATIONS

BRCA1, Breast Cancer type 1susceptibility protein; BRCT, BRCA1 C Terminus; MDC1,Mediator of DNA damage checkpoint protein 1; TopBP1, DNA Topoisomerase II Binding Protein 1; MST, Microscale Thermophoresis; RMSD, Root mean square deviation,

