## Supporting Information for "Structural features underlying the activity of benzimidazole derivatives that target phosphopeptide recognition by the tandem BRCT domain of the BRCA1 protein"

### Table S1. Thermodynamic properties of most stable water molecules observed by GIST analysis

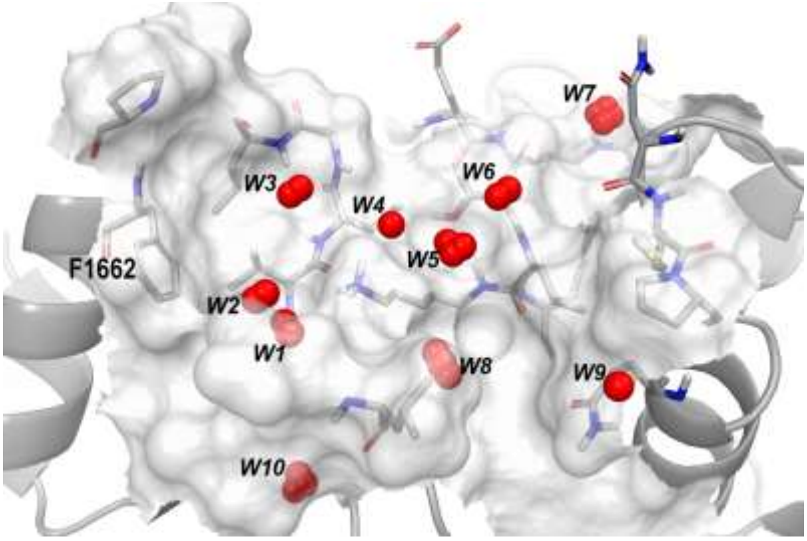


**Water grouping (red sphere) based on high density and with greater free energy change.**

| ID | $\Delta E_{sw}$ | $\Delta E_{ww}$ | $\Delta E_{total}$ | $T\Delta S^{trans}$ | $T\Delta S^{orient}$ | $T\Delta S_{total}$ | $\Delta G$ |
|---|---|---|---|---|---|---|---|
| *w1* | -14.28 | -2.33 | -16.61 | -2.540 | -2.290 | -4.830 | -11.779 |
| *w2* | -1.105 | -1.071 | -2.176 | -0.284 | -0.183 | -0.467 | -1.708 |
| *w3* | -2.473 | -1.113 | -3.586 | -0.580 | -0.304 | -0.884 | -2.702 |
| *w4* | -0.959 | -0.616 | -1.575 | -0.185 | -0.122 | -0.306 | -1.269 |
| *w5* | -3.828 | -4.034 | -7.862 | -1.600 | -0.924 | -2.524 | -5.338 |
| *w6* | -2.880 | -1.610 | -4.490 | -0.559 | -0.303 | -0.862 | -3.629 |
| *w7* | -5.997 | -2.463 | -8.460 | -1.029 | -0.605 | -1.634 | -6.827 |
| *w8* | -3.622 | -1.515 | -5.137 | -0.689 | -0.548 | -1.236 | -3.901 |
| *w9* | -1.392 | -0.287 | -1.679 | -0.166 | -0.131 | -0.298 | -1.382 |
| *w10* | -11.373 | -1.346 | -12.719 | -1.757 | -1.601 | -3.359 | -9.361 |

All values are in Kcal/mol unit

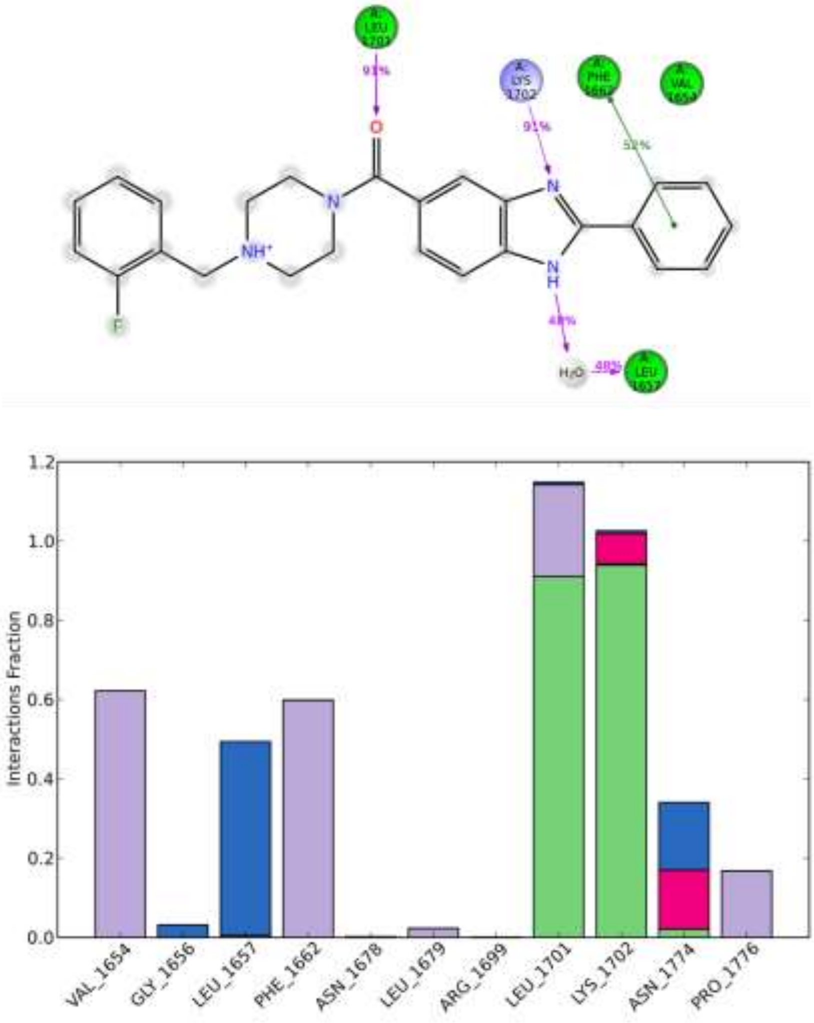


**Figure S1**. A: Two-dimensional representation of the percentage intermolecular interaction between Bractoppin and residues in the BRCA1 tBRCT binding pocket during a 5 ns MD simulation. Hydrogen bond interactions are shown in magenta, and the hydrophobic and charged pocket residues are shown in green and purple spheres, respectively. B: The histogram shows the fraction of protein- ligand contacts between Bractoppin and BRCA1 tBRCT through a 5 ns MD simulation with the pocket residues plotted on X-axis, and the interaction fraction plotted on Y-axis.

***General procedure 1:***

**Synthesis of methyl 2-phenyl-1H-benzo[d]imidazole-6-carboxylate (A):** In a vial the mixture of benzoic acid (0.367 g), methyl 3,4-diaminobezoate (0.5 g) and PPA (2 g) was heated to 170$^0$C for 3 hours, TLC (DCM : MeOH = 9:1) indicated that starting material was consumed. The reaction mixture was poured into saturated $NaHCO_3$ solution followed by extraction with ethyl acetate (20 mL x 3). The combined organic phase was washed with brine (30 mL x 2), dried with anhydrous $Na_2SO_4$, filtered and concentrated in vacuum to afford methyl 2-phenyl-1H-benzo[d]imidazole-6-carboxylate (0.12 g, crude) obtained as an off-white solid.

**Synthesis of 2-phenyl-1H-benzo[d]imidazole-6-carboxylic acid (B):** A mixture of methyl 2-phenyl-1H-benzo[d]imidazole-6-carboxylate (A) (0.1 g), concentrated HCl (7 mL), acetic acid (6 mL) was heated to 90$^o$C for 3 hours, TLC (DCM : MeOH = 9:1) indicated that starting material was consumed. The reaction mixture was neutralized by saturated $NaHCO_3$ solution followed by extraction with ethyl acetate (20 mL x 3). The combined organic phase was washed with brine (30 mL x 3), dried with anhydrous $Na_2SO_4$, filtered and concentrated in vacuum to afford methyl 2-phenyl-1H-benzo[d]imidazole-6-carboxylatic acid (0.07 g, crude) obtained as an off-white solid.

**Synthesis of (4-(2-fluorobenzyl) piperazin-1-yl) (2-phenyl-1H-benzo[d] imidazol-6-yl) methanone (Bractoppin):** To a solution of methyl 2-phenyl-1H-benzo[d]imidazole-6-carboxylic acid (0.07g) in DMF was added 1- (2-fluorobenzyl) piperazine (0.06 g) and HATU (0.17 g). Reaction mixture was cooled to 0$^0$-5$^0$C followed by the addition of DIPEA (0.1 mL) and stirred at same temperature for 2 hours, TLC (DCM-MeOH = 9:1) indicated that both starting materials were consumed. Reaction mixture was poured into water followed by extraction with ethyl acetate (10 mL x 3). The combined organic phase was washed with brine (40 mL x 2), dried with anhydrous $Na_2SO_4$, filtered and concentrated in vacuum to afford crude which was purified by flash chromatography where the product eluted at 3% MeOH in DCM to afford of (4- (2-fluorobenzyl)piperazin-1-yl) (2-phenyl-1H-benzo[d] imidazol-6-yl) methanone (0.035 g). LCMS: (M+H$^+$): 415.3. 1H NMR: DMSO-$d_6$ 400 MHz δ 13.132 (s, 1H),

8.198-8.180 (d, *J* = 7.2 Hz, 2H), 7.708-7.667 (m, 1H), 7.586-7.726 (m, 4H), 7.446-7.412 (t, *J* = 6.8 Hz, 1H), 7.341-7.324 (d, *J* = 6.8 Hz, 1H), 3.581 (s, 3H), 2.442 (s,1H), 1.225 (s,1H); HPLC Purity: 95.89%.

Compounds 2010, 2049, 2086, 2088, 2090, 2113 and 2119 were synthesized using the general procedure 1 and the the analytical data is given below.

**(2- (3,5-dimethoxyphenyl)-1H-benzo[d]imidazol-6-yl) (4- (2-fluorobenzyl) piperazin -1-yl)methanone (2010):** LCMS: (M+H$^+$): 741.22. $^1$H NMR: (MeOD, 400 MHz) δ 7.725-7.655 (br, 2H), 7.474-7.433 (td, *J* = 1.6 Hz, 1H), 7.365-7.302 (m, *J* = 2 Hz, 4H), 7.199 -7.159 (m, *J* = 0.8 Hz, 1H), 7.182-7.028 (m, *J* = 8.4 Hz, 1H), 6.682-6.671 (t, *J* = 2, 1H) , 3.909 (s, 5H), 3.810 (s, 2H), 3.685-3.661 (d, *J* = 9.6 Hz, 2H), 3.588 (br, 2H), 2.603-2.542 (br, 4H), 1.356-1.306 (m, *J* = 13.2 Hz, 1H); HPLC Purity: 99.63%.

**(S)- (4- (2-fluorobenzyl)-2-isopropylpiperazin-1-yl) (2-phenyl-1H-benzo[d] imidazole -5 -yl) methanone (2049)** LCMS: (M+H$^+$): 456.3; $^1$H NMR (DMSO-$d_6$, 400 MHz) δ 13.1654 (s, 1H), 8.195 (s, 2H), 7.566 (br s, 4H), 7.434 (br s, 1H), 7.316 (br s, 1H), 7.181 (br s, 3H), 4.225 (br s, 1H), 3.529-3.504 (br s, 3H), 2.947 (s, 2H), 2.704 (s, 1H), 2.421 (br s, 1H), 2.072 (s, 1H), 2.010 (s, 1H), 0.909-0.832 (br d, 3H), 0.669-0.505 (br d, 2H); HPLC Purity: 97.48%.

**2-(4-(6-(4-(2-fluorobenzyl)piperazine-1-carbonyl)-1H-benzo[d]imidazol-2-yl)phenoxy)acetonitrile (2088):** LCMS: (M+H+): 470.11; $^1$H NMR (DMSO-$d_6$ 400 MHz) δ 10.171 (1 1H), 7.863-7.645 (m, 4H), 7.436-7.334 (m, 3H), 7.185 (d, *J* = 4.4 Hz, 2H), 6.999 (d, *J* = 8.0 Hz, 2H), 5.617 (d, 2H), 3.844-3.329 (m, 6H), 2.646-2.547 (m, 4H); HPLC Purity: 99.26%.

**(4-(2-fluorobenzyl)piperazin-1-yl)(2-(5-methyl-1,3,4-oxadiazol-2-yl)-1H-benzo[d] imidazol-5-yl)methanone (2086):** LCMS: (M+H$^+$): 421.33; $^1$H NMR (DMSO-$d_6$ 400 MHz) (80$^0$C), δ 13.640 (br s, 1H), 7.630-7.780 (m, 2H), 7.439 (t, J=7.6 Hz, 1H), 7.302-7.365 (m, 2H), 7.120-7.200 (m, 2H), 3.620 (s, 2H), 3.500-3.590 (m, 4H), 2.669 (s, 3H), 2.450-3.450 (m, 4H); HPLC Purity: 96.80%.

**(4-(2-fluorobenzyl)piperazin-1-yl)(2-(4-methoxyphenyl)-1H-benzo[d]imidazol-5-yl)methanone (2090):** LCMS: (M+H$^+$): 445.5; $^1$H NMR (MeOD 400 MHz) δ 8.048-8.077 (m, 2H), 7.620-7.690 (m, 2H), 7.732-7.7.447 (m,

1H), 7.303-7.359 (m, 2H), 7.104-7.199 (m, 4H), 3.909 (s, 3H), 3.570-3.870 (m, 6H), 2.491-2.680 (m, 4H); HPLC Purity: 95.69%.

**2-(5-(4-(2-fluorobenzyl)piperazine-1-carbonyl)-1H-benzo[d]imidazol-2-yl)acetonitrile(2091):** LCMS: (M+H$^+$): 378.4; $^1$H NMR (MeOD, 400 MHz) δ 7.580.7.730 (m, 2H), 7.434-7.472 (m, 1H), 7.314-7.366 (m, 2H), 7.182 (t, *J* = 7.6 Hz, 1H), 7.109 (t, *J* = 1.6 Hz, 1H). 3.50-4.00 (m, 8H), 2.500-2.700 (m, 4H); HPLC Purity: 95.65%.

**(S)-2-(5-(4-(2-fluorobenzyl)-2-isopropylpiperazine-1-carbonyl)-1H-benzo[d]imidazol-2-yl)acetonitrile (2076):** LCMS: (M+H$^+$): 420.26 $^1$H NMR (DMSO-$d_6$, 400 MHz) δ 12.726-12.796 (d, *J* = 28 Hz, 1H), 7.321-7.633 (br, 4H), 7.184 (br, 3H), 4.423 (s, 2H), 4.214 (br, 1H), 3.526 (br, 3H), 2.944 (br, 2H), 2.677 (br, 1H), 2.406 (br, 1H), 2.191 (br, 1H), 2.004 (br, 1H), 0.898-0.836 (d, 4H), 0.648 (bs, 1H), 0.498 (bs, 1H); HPLC Purity: 95.62%.

**(S)-(4-(2-fluorobenzyl)-2-isopropylpiperazin-1-yl)(2-(3-(4-methoxyphenoxy)propyl)-1H-benzo[d]imidazol-5-yl)methanone (2077):** LCMS: (M+H$^+$): 545.4; $^1$H NMR (DMSO-$d_6$, 400 MHz) δ 12.435-12.395 (d, *J* = 16 Hz, 1H), 7.452-7.417 (m, 2H), 7.354-7.302 (bm, 1H), 7.202-7.149 (m, 2H), 7.107 (bs, 1H), 6.877-6.824 (bm, 3H), 4.217-4.196 (d, *J* = 8.4 Hz, 1H), 4.019-3.989 (t, *J* = 6 Hz, 2H), 3.686 (s, 2H), 3.526-3.464 (bm, 2H), 3.341 (bs, 1H), 3.006-2.942 (m, 2H), 2.704-2.675 (d, *J* = 11.6 Hz, 1H), 2.403 (bs, 1H), 2.239-2.171 (bm, 2H), 2.026-1.963 (bm, 1H), 0.892-0.834 (d, *J* = 23.2 Hz 3H), 0.502 (s, 1H); HPLC Purity: 97.89%.

**(4-(2-fluorobenzyl)piperazin-1-yl)(2-(3-(4-methoxyphenoxy)propyl)-1H-benzo[d]imidazol-5-yl)methanone (2113):** LCMS: (M+H$^+$): 503.23; $^1$H NMR (MeOD, 400 MHz) δ 7.578-7.595 (m, 2H), 7.435-7.469 (m, 1H), 7.285-7.365 (m, 2H), 7.183 (t, *J* = 6.8 Hz, 1H), 7.108 (t, *J* = 9.6 Hz, 1H), 6.770-6.850 (m, 4H), 4.018 (t, *J* = 6.0 Hz, 2H), 3.50-3.90 (m, 9H), 3.130 (t, *J* = 7.6 Hz, 2H), 2.500-2.700 (m, 4H), 2.292-2.328 (m, 2H); HPLC Purity: 98.62%.

**(4-(2-fluorobenzyl)piperazin-1-yl)(2-(4-(2-methoxyethoxy)phenyl)-1H-benzo[d]imidazol-5-yl)methanone (2119):** LCMS: (M+H$^+$): 488.9: $^1$H NMR (DMSO-$d_6$, 400 MHz) δ 12.96 (1H), 8.117 (d, *J* = 8.4 Hz, 2H), 7.578-7.595 (m, 2H), 7.419-7.453 (m, 1H), 7.333-7.347 (m, 1H), 7.183-7.250 (m, 5H), 4.196 (m, 2H),

3.699 (m, 2H), 3.4-3.6 (m, 5H), 3.3-3.4(m, 3H), 2.350-2.500 (m, 4H); HPLC Purity: 97.04%.

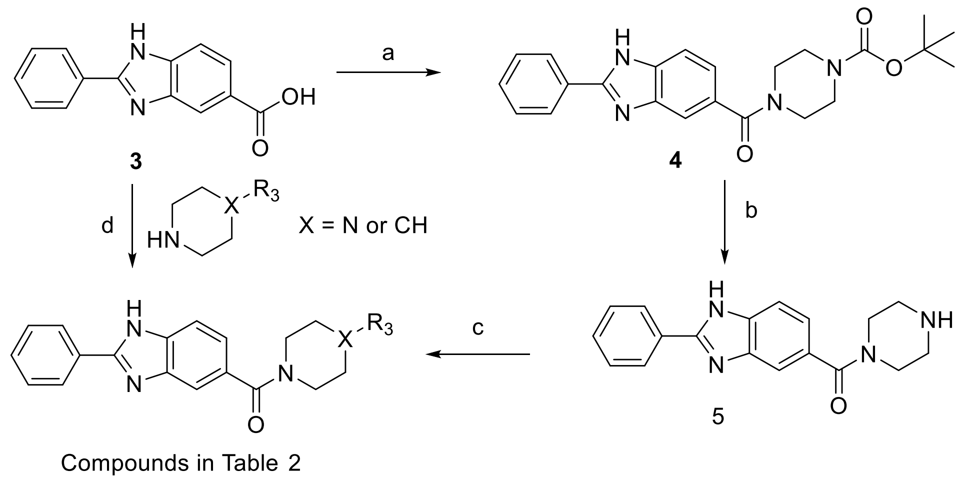


**Scheme 2:** Reagents and conditions: (a) tert-butyl piperazine-1-carboxylate, HATU, DIPEA, DMF; (b) 4N HCl.Dioxane; (c) $Na(CH_3COO)_3BH$, AcOH, DCM; (d) EDC, HOBt, DMF.

***General procedure 2:***

***Synthesis of Intermediate-4:***

To a solution of Intermediate-3 (2.0 g, 1.5 mmol, 1.0 eq) in DMF (10 mL) were added HATU (0.684 g, 1.8 mmol, 1.2 eq) and DIPEA (0.76 mL, 4.5 mmol, 3.0 eq) at 0 °C. the reaction mixture was stirred for 30 min at room temperature. tert-butyl piperazine-1-carboxylate (0.286 g, 1.5 mmol, 1.0 eq) was added in to the reaction mixture and stirred room temperature for 1.5 h, TLC ($CHCl_3$.MeOH = 9:1) indicated the starting material was consumed. Reaction mixture was poured into ice cold water (70 mL) and extracted with ethyl acetate (50 mL*3). The combined organic phase was washed with brine (100 mL*2), dried over anhydrous $Na_2SO_4$, filtered and concentrated under reduced pressure to afford Intermediate-4 (2.0 g,) as a pale yellow solid.

***Synthesis of Intermediate-5***

To a solution of Intermediate-4 (2.0 g, 4.91mmol, 1.0 eq) in 1, 4-dioxane was added 4N HCl in 1, 4-dioxane (6 mL) at 0 °C. The reaction mixture was stirred at room temperature for 2 hour, TLC ($CHCl_3$.MeOH = 9:1) indicated the starting material was consumed. The reaction mixture was concentrated in vacuum to afford Intermediate-5 (1.5 g) was obtained as, light brown solid (as HCL salt).

***Synthesis of* (4-(2-methoxybenzyl)piperazin-1-yl)(2-phenyl-1H-benzo[d]imidazol-5-yl)methanone (2096):** To a solution of Intermediate-5 (0.25 g, 0.730 mmol, 1.0 eq) and 2-methoxybenzaldehyde (1.0 g, 0.803 mmol, 1.1 eq) in DCM (10 mL) were added TEA (0.113 g, 1.125 mmol, 1.5 eq) and acetic acid (5 drops) at room temperature. The reaction mixture was stirred at room temperature for 1 h. Sodium triacetoxyborohydride (1.0 g, 2.19 mmol, 3.0 eq) was added in to the reaction mixture and stirred for 20 h, TLC ($CHCl_3$.MeOH = 9:1) indicated the starting material was consumed. Reaction mixture was poured into ice cold water and extracted with ethyl acetate (50 mL*3). The combined organic phase was washed with brine (25 mL*2), dried with anhydrous $Na_2SO_4$, filtered and concentrated in vacuum to afford the crude which was purified by prep-HPLC purification using $NH_4HCO_3$ as buffer to afford 2096 (0.15 g) as off white solid. LCMS: (M+H$^+$): 427.5; $^1$H NMR **(**DMSO-$d_6$, 400 MHz) δ 13.122 (s, 1H), 8.193 (d, J = 8.4 Hz, 2H), 7.673-7.713 (m, 1H), 7.508-7.591 (m, 4H), 7.330 (d,J=7.2, 1H), 7.209-7.279 (m, 2H), 6.911-6.994 (m, 2H), 3.778 (s, 3H), 3.510-3.610 (m, 6H), 2.410-2.490 (m, 4H); HPLC Purity: 99.84%.

Compounds 2103 and 2104 were synthesized using the general procedure 2 and the analytical data is given below.

**(4-(3-morpholinobenzyl)piperazin-1-yl)(2-phenyl-1H-benzo[d]imidazol-6-yl)methanone (2103):** LCMS: (M+H$^+$): 482.6; $^1$H NMR (DMSO-$d_6$, 400 MHz) δ 13.168 (s, 1H), 8.198 (d, *J* = 7.2 Hz, 2H), 7.686-7.712 (m, 1H), 7.507-7.594 (m, 4H), 7.174-7.282 (m, 2H), 6.784-6.920 (m, 3H), 3.736 (t, *J* = 9.6 Hz, 4H), 3.310-3.670 (m, 6H), 3.096 (t, *J* = 9.2 Hz, 4H), 2.410-2.510 (m, 4H); HPLC Purity: 95.09%.

**(4-((1H-pyrazol-5-yl)methyl)piperazin-1-yl)(2-phenyl-1H-benzo[d]imidazole-6-yl)methanone (2104):** LCMS: (M+H$^+$): 387.5; $^1$H NMR (DMSO-$d_6$, 400 MHz): δ 13.140 (s, 1H), 12.603 (br s, 1H), 8.205-8.185 (d, *J* = 8.0 Hz, 2H), 7.510-7.750 (m, 5H), 7.210-7.90 (m, 1H), 6.165 (s, 1H), 5.769(s, 1H), 3.450-3.650 (m, 6H), 2.350-2.500 (m, 4H); HPLC Purity: 999.26%.

***General Procedure 3:***

**Synthesis of (4-methylpiperazin-1-yl)(2-phenyl-1H-benzo[d]imidazol-5-yl)methanone (2048):** To a solution of Intermediate-3 (0.15 g, 0.63 mmol, 1.0 eq) in DMF (10 mL) was added EDC.HCl (0.13 g, 0.69 mmol, 1.1 eq) and HOBt (0.04 g, 0.31 mmol, 0.5 eq) and stirred at room temperature for 30 minutes. To this Nmethyl piperazine (0.06 g, 0.63 mmol, 1.0 eq) and DIPEA (0.3 mL, 1.89 mmol, 3.0 eq) was charged. The mixture was stirred at room temperature for 18 hours, TLC ($CHCl_3$:MeOH = 9:1) indicated the starting material was consumed. Reaction mixture was poured into water followed by extraction with ethyl acetate (30 mL*3). The combined organic phase was washed with brine (40 mL*2), dried with anhydrous $Na_2SO_4$, filtered and concentrated in vacuum to afford the crude which was purified by flash chromatography where the product eluted at 3.9% MeOH in chloroform followed by trituration with n-pentane to afford 2048 (0.067g) as yellow solid. LCMS: ($M+H^+$): 321.4; $^1H$ NMR: DMSO-$d_6$ 400 MHz) δ 13.137 (s, 1H), 8.203-8.185 (d, *J* = 7.6Hz, 2H), 7.716-7.675 (t, *J* = 8.0 Hz, 2H), 7.594-7.523 (m, 4H), 7.277-7.207(m, 1H), 3.533 (broad s, 4H), 2.337 (broad s, 4H), 2.210 (d, 3H); HPLC Purity: 99.81%.

Compounds 2098 and 2099 were synthesized using the general procedure 3 and the analytical data is given below.

**(4-benzylpiperazin-1-yl)(2-phenyl-1H-benzo[d]imidazol-6-yl)methanone**

**(4-phenethylpiperazin-1-yl)(2-phenyl-1H-benzo[d]imidazol-6-yl)methanone (2098):** LCMS: ($M+H^+$): 411.5; $^1H$ NMR (DMSO-$d_6$, 400 MHz): (80$^0$C) δ 9.610 (br s, 1H), 8.188-8.212 (m, 2H), 7.756 (s, 1H), 7.687 (d, *J* = 8.4 Hz, 1H), 7.555-7.615 (m, 3H), 7.357-7.394 (m, 3H), 7.291-7.312 (m, 2H), 4.100-4.400 (m, 4H), 3.200-3.700 (m , 6H), 3.033-3.075 (m, 2H); HPLC Purity: 97.87%.

**(4-(2-fluorobenzyl)piperidin-1-yl)(2-phenyl-1H-benzo[d]imidazol-6-yl)methanone (2099):** LCMS: ($M+H^+$): 414.5; $^1H$ NMR (DMSO-$d_6$, 400 MHz): δ 13.075 (br s, 1H), 8.194 (d, *J* = 6.8 Hz, 2H), 7.506-7.595 (m, 5H), 7.235-7.309 (m, 3H), 7.117-7.170 (m, 2H), 4.450-4.520 (m, 1H), 3.620-3.810 (m , 1H), 2.720-2.890 (m, 2H) 2.605 (d, *J* = 7.2 Hz, 2H), 1.500-1.890 (m, 3H), 1.150-1.300 (m , 2H); HPLC Purity: 99.66%.
